# BZIP Transcription Factors Modulate DNA Supercoiling Transitions

**DOI:** 10.1101/2019.12.13.875146

**Authors:** Johanna Hörberg, Anna Reymer

## Abstract

Torsional stress on DNA, introduced by molecular motors, constitutes an important regulatory mechanism of transcriptional control. Torsional stress can modulate specific binding of transcription factors to DNA and introduce local conformational changes that facilitate the opening of promoters and nucleosome remodeling. Using all-atom microsecond scale molecular dynamics simulations together with a torsional restraint that controls the total helical twist of a DNA fragment, we addressed the impact of torsional stress on DNA complexation with a human BZIP transcription factor, MafB. We gradually over- and underwind DNA alone and in complex with MafB by 5° per dinucleotide step, monitoring the evolution of the protein-DNA contacts at different degrees of torsional strain. Our computations show that MafB changes the DNA sequence-specific response to torsional stress. The dinucleotide steps that are susceptible to absorb most of the torsional stress become more torsionally rigid, as they are involved in the protein-DNA contacts. Also, the protein undergoes substantial conformational changes to follow the stress-induced DNA deformation, but mostly maintains the specific contacts with DNA. This results in a significant asymmetric increase of free energy of DNA twisting transitions, relative to free DNA, where overtwisting is more energetically unfavorable. Our data suggest that MafB could act as a torsional stress insulator, modulating the propagation of torsional stress along the chromatin fiber, which might promote cooperative binding of other transcription factors.

---

Torsional restraints on DNA, referred to as DNA supercoiling, constantly change during the life of the cell, and regulate transcriptional control on many levels.^1–5^ DNA supercoiling represents a sum of writhe and twist – the two interchangeable variables. DNA writhing generally dominates supercoiling changes on a larger scale through the formation of loops and knots, while DNA twisting occurs when shorter DNA fragments, up to ~100 base pairs (b.p.), experience changes in torsional restraints. The net state of genomic DNA is neutral, but regions of positive and negative supercoiling can exist locally, created by RNA polymerases that expose DNA to torsional stress.^3,5^ This introduces DNA undertwisting (negative supercoiling) upstream and overtwisting (positive supercoiling) downstream of a transcribed gene.

Torsional stress can propagate along DNA, modulating transcription of near-located genes^1,5^ by altering the stability of nucleosomes and other protein-DNA complexes,^3,4,6,7^ changing the accessibility of the genetic code. The ranges and speeds of torsional stress propagation depend on the underlying nucleotide sequence.^1^ Computational experiments confirm: DNA responds to torsional stress in a heterogeneous and sequence-dependent manner.^8,9^ Certain dinucleotide steps, mainly pyrimidine-purine (YpR) but also purine-purine (RpR), in specific sequence environments, absorb a large part of DNA over- and undertwisting, while the rest of the molecule preserves its relaxed B-like conformation. The torsional plasticity of these dinucleotides is founded in the polymorphic nature of the DNA backbone.^10–12^ When absorbing torsional stress, these dinucleotide steps favor respectively low (DNA-underwinding) or high (DNA-overwinding) twist states, which are separated by about 20°. The twist transitions are coupled with significant changes in other helical parameters, such as shift and slide. We hypothesize that these dinucleotide steps are potential ‘hot spots’ for transcriptional control, as they can regulate supercoiling transitions, the deformability of DNA, and specific binding by transcription factors.

Transcription factors (TFs), while operating in the large excess of non-specific DNA, must unmistakably bind their corresponding DNA targets to correctly initiate transcription reactions. The specific binding of TFs is usually considered in terms of three mechanisms. (1) The ‘direct readout’ which involves the formation of specific hydrogen bonds and hydrophobic interactions between DNA bases and protein amino acids.^13–15^ (2) The ‘indirect readout’ of the DNA shape.^16–18^ (3) The water-mediated interactions between DNA and protein.^19–21^ The three recognition modes can contribute differently to the specificity of TF-DNA binding, depending on the type of TF and the recognized DNA sequence. Irrespective of the dominating recognition mechanism, torsional stress passing through the genome will change the geometry of the DNA helix, and potentially alter the stability of a TF-DNA complex. Presence of a protein will likely affect the free energy of DNA torsional stress propagation, potentially regulating transcription of nearby genes. Despite being central for eukaryotic transcriptional control, these mechanistic aspects of TFs-DNA interactions are far from being understood in detail.

Motivated by the scarcity of mechanistic studies, we conduct a computational experiment where we apply torsional stress to DNA in complex with MafB^22–24^, a member of the BZIP family of human TFs. We perform all-atom umbrella sampling simulations using a torsional restraint that controls the total twist of a DNA molecule.^8^ The restraint does not restrict any other degrees of freedom of DNA and can be applied to DNA alone and in complex with proteins while running molecular dynamics simulations. We observe that MafB locks the most torsionally flexible dinucleotide steps from oscillating between low- and high-twist substates, through forming specific contacts with the bases. As a result, DNA becomes asymmetrically more rigid, with DNA overtwisting being more energetically unfavorable.

We simulate two systems – the MafB-DNA complex (PDB ID: 4AUW)^22^ and free DNA. In both systems, we use the following DNA sequence ‘GGTAAT **TGCTGACGTCAGCA**TTATGG’, with the MafB-response element (MARE) in bold. To recognize MARE-DNA, MafB dimer utilizes the direct readout mechanism, where a six-residues motif (**R**xxx**N**xx**Y-A**xx**CR**) of each monomer forms specific contacts via the major groove to the MARE-half site – TGCTGAC. The structural details of MafB-DNA recognition are described elsewhere^25^ (see also Figures S1, S2). We apply the twist restraint to the 14 b.p. MARE-region, gradually increasing (overwinding) or decreasing (underwinding) the total twist by 7°(0.5°/b.p.), starting from the relaxed state to a maximum of ±5°/b.p. We perform a cascade umbrella sampling, with 0.5 microsecond simulation time per window, 21 windows in total, to obtain the potential of mean force (PMF) of DNA twisting free energy as a function of average b.p. twist. We use a force constant *k_tw_* of 0.06 kcal/mol×degrees^2^, which provides the desired torsional strain without introducing any structural anomalies. For details of the simulation protocol see Supporting Information.

The PMF profiles (Figure 1) show the energy cost for DNA twisting asymmetrically increases in the presence of MafB, where overtwisting becomes noticeably more unfavorable. An additional 2.9 kcal/mol has to be paid to overwind complexed DNA by 5°/b.p. (70° in total) with respect to free DNA. In contrast, for the same degree of underwinding the energy difference is much smaller, 1.1 kcal/mol. The derived torsional force constants of 0.06 and 0.11 kcal/mol×degrees^2^, and the torsional moduli of 442 and 853 pN×nm^2^, for free and complexed MARE-region, correspondingly, indicate that DNA in complex with the transcription factor becomes nearly twice more rigid.

**Figure 1.**
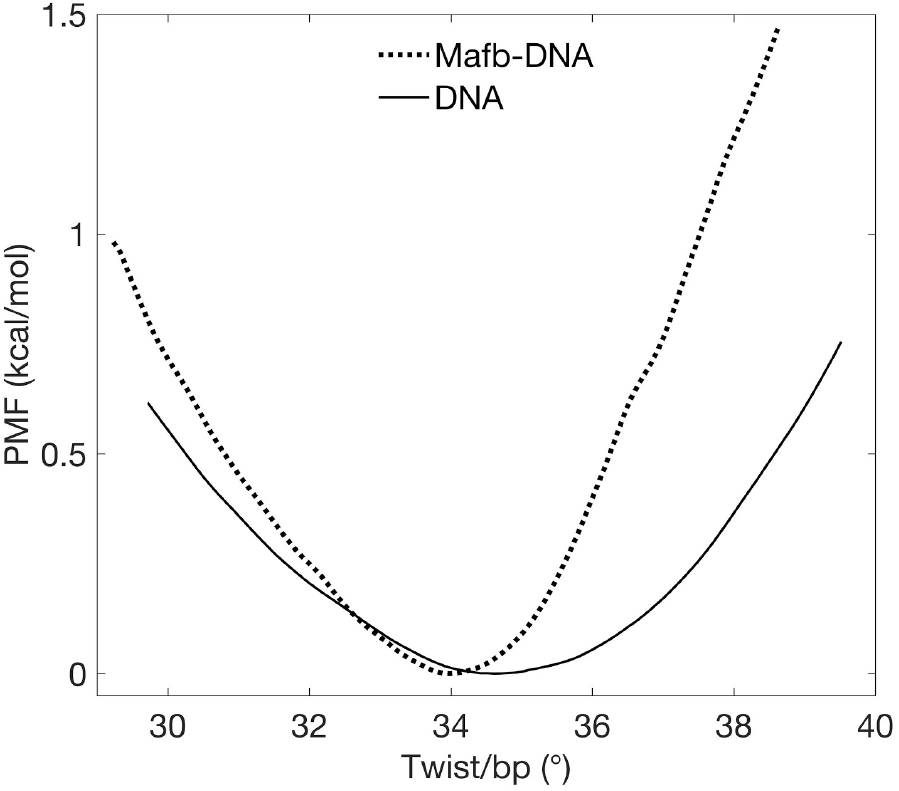
PMF profiles with respect to average twist per base pair step in free and complexed MARE-DNA.

To explain the mechanism of the induced rigidity by MafB complexation, we analyze the response of the individual b.p. steps to torsional stress (Figure 2). In accordance with our previous studies^8,9^, the TpG and CpA steps of C-TpG-A and T-CpA-G motifs exhibit major torsional flexibility in free DNA, effectively absorbing both negative and positive torsional stress. The two tetranucleotide motifs are symmetrical, each belonging to the MARE-half site. In the presence of MafB, however, the TpG and CpA steps become rigid. Twist distributions for the steps, which show twist bimodality in free relaxed DNA, exhibit a high twist state in the protein environment. During DNA overwinding, these steps remain passive, as they are unable to increase their twist further. The contribution of the TpG and CpA steps to efficient DNA underwinding is also limited. Instead, other b.p. steps that are less flexible in free DNA are forced to modify their twist, resulting in the increased energy cost for DNA twisting. We exclude the first and the last b.p. step of the restrained region from the analysis since the variation of their twist values may result also from the boundary effects. For the comparison of twist distributions for the restrained MARE-region for the relaxed, overwound (+4.5°), and underwound (−4.5°) states see figure S4.

**Figure 2.**
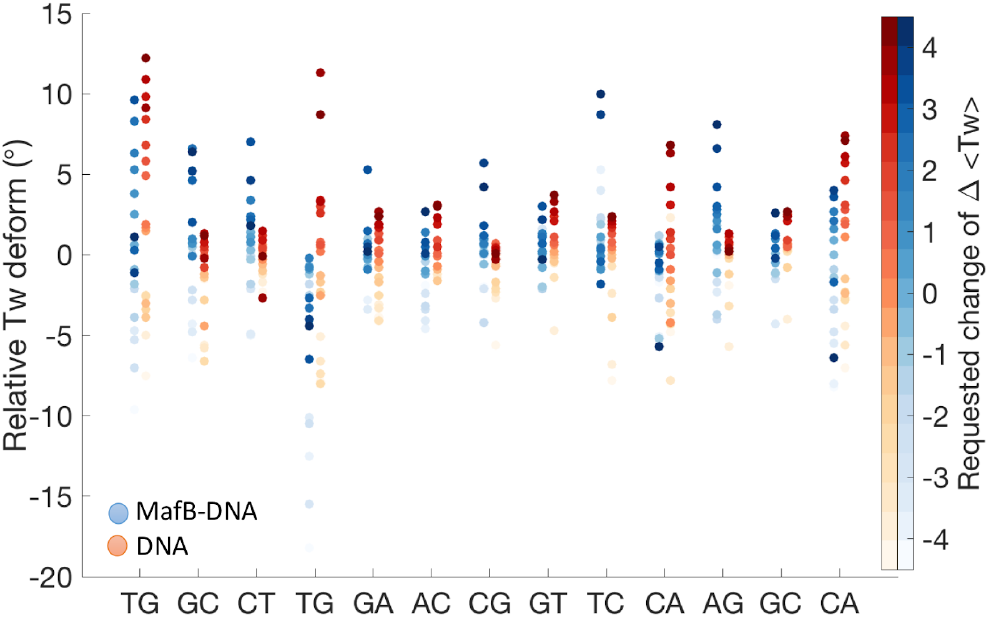
Changes of twist angles for the restrained MARE-region in free and complexed DNA as a function of the requested average change of twist per base pair step, indicated by a colorbar to the right.

We consider further the protein role in the modified DNA sequence-specific response to torsional stress. Upon association with the MARE-region, MafB induces local structural adjustments of the double helix. In particular, the TpG step, which hydrophobically interacts with Tyr251, Ala252, and Cys255 of RxxNxx**YA**xx**C**R-motif, exhibits a negative shift – the G-C b.p. is effectively pushed towards the DNA minor groove (Figure S5). The same behavior is observed for the symmetric CpA step in the other MARE-half site. This allows the protein to have tighter hydrophobic contacts with the T-A b.p. of the MARE-half sites. Since the helical parameters shift and twist are structurally coupled via the BI/BII backbone conformational transitions,^12,8,9^ the protein-induced negative shift locks the TpG and CpA dinucleotides in a high twist substate.

We monitor the evolution of the MafB-DNA contacts network to estimate the structural impact of changing torsional stress on the protein-DNA complex. Since the MafB-MARE recognition process follows the direct readout mechanism, we hypothesize that the number of contacts represents the complex stability. We characterize the protein-DNA interactions by pairs of residues, dividing the contacts into ‘specific’, formed between the protein side chains and DNA bases, and ‘non-specific’, formed with at least one of the molecules’ backbones. For each pair of protein-DNA residues, we sum up all hydrogen bonds, salt bridges, and hydrophobic (apolar) interactions. The contribution of a single bond of each type is set to 1 (Figure S7), for simplicity, since the energy cost of every type of bond vary greatly depending on the nature of atoms involved, the bond geometry, and the surrounding environment. The time series of MafB-DNA interactions allow the construction of dynamic contact maps for specific (Figure 3) and non-specific contacts (Figure S8), characterizing the stability and the binding specificity of the MafB-DNA complex at different degrees of positive and negative torsional stress.

**Figure 3.**
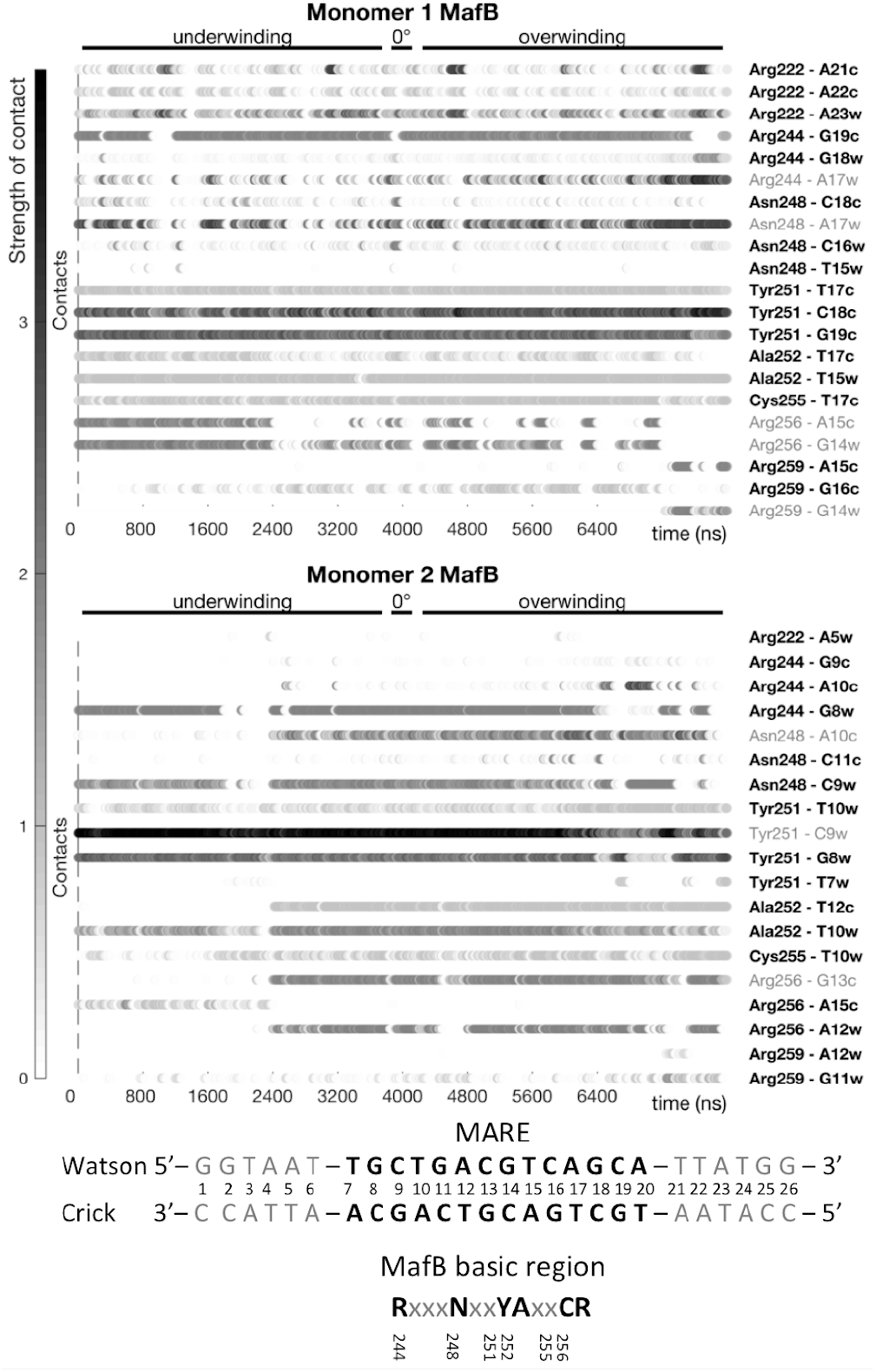
Dynamic interactions maps illustrate specific MafB-DNA contacts at different degrees of torsional stress. The contacts between pairs of residues are characterized by strength and occurrence. Torsional stress denoted as ‘underwinding’ represents changes from −5°/b.p. to −0.5°/b.p.; and ‘overwinding’ – from 0.5°/b.p. to 5.0°/b.p. Text in bold shows contacts that change insignificantly (change in contact strength < 1) with changing torsional stress.

The dynamic contact maps show that despite torsional stress, MafB maintains the majority of the specific contacts with the MARE-binding site. However, we observe some changes in the intermolecular contacts network during DNA underwinding. The MARE-half site on Watson-strand absorbs most of the negative torsional stress, making the T12c and G13c bases inaccessible for MafB monomer 2. This leads to the loss of specific contacts: Ala252(2)-T12c and Arg256(2)-G13c, at higher levels of underwinding < −2.5°/b.p. Instead, a compensating contact Arg256(2)-A15c is formed. The MARE-half site on Crick strand exhibits a tighter interaction with MafB monomer 1, which effectively makes it more torsionally rigid. Furthermore, DNA underwinding stabilizes two specific contacts, Arg256(1)-G14w and Arg256(1)-A15c. The most noticeable change during extreme DNA overwinding, > 4°/b.p., is formation of a specific contact, Arg259(1)-G14w. The dynamic contact maps for the non-specific contacts (Figure S8) mirror the trends of the specific contacts, namely, the mentioned arginine residues gain or lose, respectively, contacts with the DNA backbone. We also observe that to maintain the contacts with torsionally stressed DNA, MafB modifies the structure of its DNA binding domains. At the higher degree of underwinding, < −4.0°/b.p., the long alpha-helices appear to bend away from DNA, while for the higher degree of overwinding, > 4.0°/b.p., the helices bend towards DNA (Figure S9).

The observed cooperative mechanism, when one protein monomer compensates for the loss of contacts between another monomer and torsionally stressed DNA, we propose, is characteristic for BZIPs. The long alpha-helical coiled-coil DNA binding domains of BZIPs can adjust to the torsionally modified geometry of the double helix to maintain stable contacts with DNA. Our observations suggest that BZIPs may act as topological insulators, hindering the propagation of torsional stress along the chromatin fiber. The stable binding of BZIP factors, we hypothesize, may initiate a formation of enhanceosomes, acting as pioneer factors that inhibit chromatin compaction.

## ACKNOWLEDGMENT

This work was supported by Swedish Foundation for Strategic Research SSF Grant [ITM170431] and Hasselblad Foundation Prize to A. R. The authors thank Dr. Elisa Frezza and Dr. Alexey Voronov for helpful discussions, and Swedish National Infrastructure for Computing (SNIC) for the generous provision of computing resources.

## Supporting Information

Details of simulation protocols and analysis, additional figures, and movie illustrating the structural deformation of the protein.

