## Supporting information for "BZIP Transcription Factors Modulate DNA Supercoiling Transitions"

---

### DNA Systems

We studied two systems: the MafB-DNA complex (PDB ID: 4AUW)[1] and free DNA. Both systems contained a DNA 26-mer oligomer: GGTAAT**TGCTGACGTCAGCA**TTATGG. For each system, the twist restraint[2] was applied to the central MARE region between b.p. 7 and b.p. 20; 13 b.p. steps in total. Starting from fully relaxed state, the total twist of the MARE region was gradually increased and decreased in steps of 0.5°/b.p step (±6° in total), until a maximum overwound and underwound state of 5°/b.p. step was reached (±65° in total). The final structure from every window was used as starting point for the adjacent window, following a so-called umbrella sampling scheme[3], which has proven to provide good convergence.

### Molecular Dynamics Simulation Protocol

Structural preparation of the MafB-DNA complex and free DNA systems were done in the modeling program JUMNA[4]. Molecular dynamics (MD) simulations and cascade umbrella sampling were performed using the MD engine GROMACS v5.1[5]. In addition, the restrained MD simulations were carried out with the help of a torsional restraint[2] that control the total helical twist of a DNA segment, which has been implemented via PLUMED v2.2[6]. The force constant ($K_{tw}$) was set to 0.06 kcal $mol^{-1}$ $degrees^{-2}$, which enables the desired torsional strain but without inducing any structural artifacts. All simulations were performed using a combination of AMBER 14SB[7] and Parmbc1[8] force fields to treat the protein and DNA respectively.

The MafB-DNA complex and free DNA oligomer were separately solvated in truncated octahedron periodic boxes by SPC/E water molecules[9] with a buffer distance of 15 Å to the walls, and subsequently neutralized by K+ counterions. Additional K+ and Cl- ions were then added to reach a physiological salt-concentration of 150 mM. Applying periodic boundary conditions, each system was subjected to energy minimization with 5000 steps of steepest descent, followed by a 500 ps equilibration-run at constant volume, raising the temperature to 300 K. Simulations were then carried out at constant pressure and temperature (1 atm and 300 K), where the temperature was controlled by a weak-coupling thermostat[10] with a coupling constant of 0.2 ps and the pressure was controlled by an isotropic Paririnello-Rahman barostat[11] with a coupling constant of 2 ps. All bonds involving hydrogen atoms were constrained with the LINCS algorithm[12], allowing to set the time step to 2 fs. Electrostatic interactions were treated with the Particle Mesh Ewald[13] summation method using a short-range cutoff of 10 Å. The van-der-Waals forces were also truncated at 10 Å with added long-range corrections. The neighbor pair list for nonbonded interactions was updated every 20th step through the Verlet cutoff scheme[14]. Center of mass movement was removed every 0.2 ps to eliminate translational kinetic energy[15].

The initial 100 ns run of NPT MD, considered as equilibrations, was followed by a production run of 0.5 µs. Following unrestrained MD simulations, the cascade umbrella sampling was performed with 0.5 μs sampling time per window to allow sufficient convergence of DNA conformational substates and ion populations[16]. The Weighted Histogram Analysis Method (WHAM)[17], implemented in PLUMED was used to derive the potential of mean force (PMF) with respect to DNA twisting. Discarding the initial 100 ns as equilibration, two blocks containing 200 ns from each sampling window were created, and WHAM method was applied to test for convergence. The procedure showed negligible deviation (< 5%) of the provided PMF profiles. The total simulation time was 11 μs for each system.

### Conformational Analysis

The recorded trajectories were processed in CPPTRAJ[18] program from AMBERTOOLS 16 software package. Subsequently, Curves+ and Canal[19] programs were used to derive the helical parameters, backbone torsional angles and groove geometry parameters for each trajectory snapshot extracted at 1 ps intervals. This allowed for complete time-dependent information on the impact of the MafB-DNA complexation on DNA response to torsional stress.

### Contact-Network Analysis

Analysis of the protein-DNA contacts network was performed using CPPTRAJ[18] program from AMBERTOOLS 16 software package, for each trajectory snapshot extracted at 1 ps intervals. We exclude the protein-DNA contacts that are present for less than 10% of the time in one umbrella window. We characterize the protein-DNA interactions by the pairs of residues, dividing the contacts into 'specific', formed between the protein side chains and DNA bases, and 'non-specific', formed with at least one of the molecules' backbones. For each pair of protein-DNA residues we sum up all hydrogen bonds, salt bridges, and hydrophobic (apolar) interactions (Figure S7). The contribution of each type of contact is set to 1, for simplicity, since the energy cost of hydrogen bonds, salt bridges, and hydrophobic interactions varies greatly, depending on the nature of the atoms involved, the bond geometry and the surrounding environment. The limit of a direct interaction was set up to be ≤ 4 Å for a hydrogen bond between the relevant heavy atoms, and the angle limit was set up to ≥ 135° at the intervening hydrogen atom. For a salt bridge interaction, the limit was set up to 4.0 Å between the end-group nitrogen of lysine and arginine, and DNA phosphate group. A hydrophobic interaction was defined as a "dry" contact ≤ 6Å between the centers of mass of hydrophobic residues (Ala, Ile, Leu, Met, Phe, Trp, and Cys) and DNA bases. The time series of MafB-DNA interactions allow construction of the dynamic contacts maps for specific and non-specific contacts, characterizing the stability and the binding specificity of the MafB-DNA complex at various degrees of positive and negative torsional stress.

### Additional Information

MatLab software was used for post-processing and plotting of all data. USCF Chimera[20] was used for creating molecular graphics.

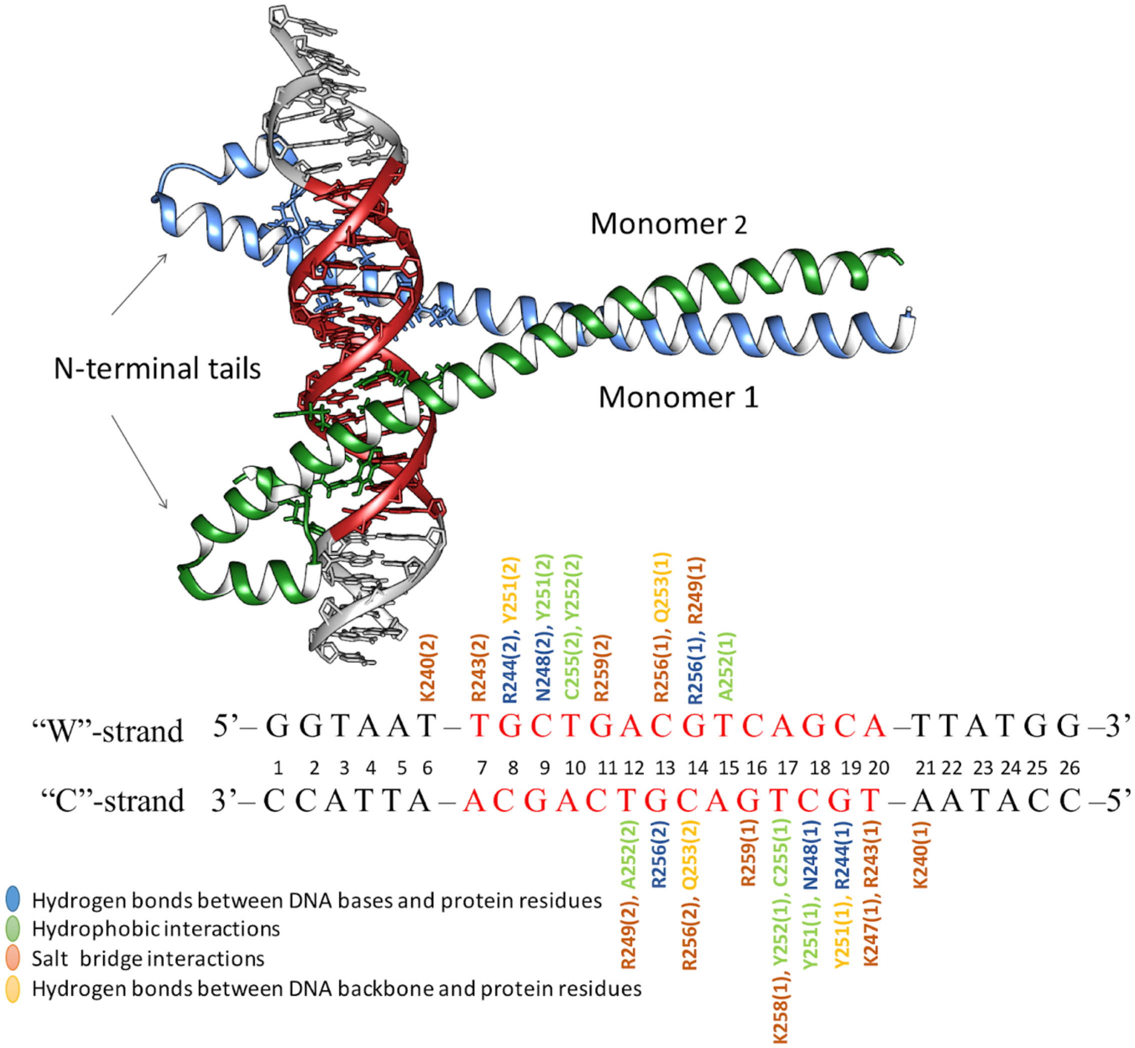


***Figure S1.*** Crystal structure of MafB-DNA complex (PDB ID: 4AUW)1, containing the MARE region denoted in red. The contacts between MafB and DNA that are present in the complex are highlighted.

MafB-DNA Contacts Involving the Conserved Six Residue Motif

**R**xxx**N**xx**YA**xx**CR**

**Specific**
Hydrophobic
Nonspecific

| Prot. Res. | Monomer 1 | Monomer 2 |
|---|---|---|
| Arg244 | **G19c** | **G8w** |
| Asn248 | **C18c** | **C9w** |
| Tyr251* | G19c<br>C18c<br>T17c | G8w<br>C9w<br>T10w |
| Ala252 | T15w | T12c |
| Cys255 | T17c | T10w |
| Arg256 | C13w<br>**G14w** | **G13w**<br>C14w |

*: Tyr251 stabilises the orientation of Arg244 and Asn248.

"W"-strand 5'– G G T A A T – T G C T G A C G T C A G C A –T T A T G G –3'
1 2 3 4 5 6 7 8 9 10 11 12 13 14 15 16 17 18 19 20 21 22 23 24 25 26
"C"-strand 3'– C C A T T A – A C G A C T G C A G T C G T – A A T A C C –5'

***Figure S2.*** Intermolecular interactions between the DNA-recognizing motif of MafB monomers and DNA, present in the crystal structure.

A

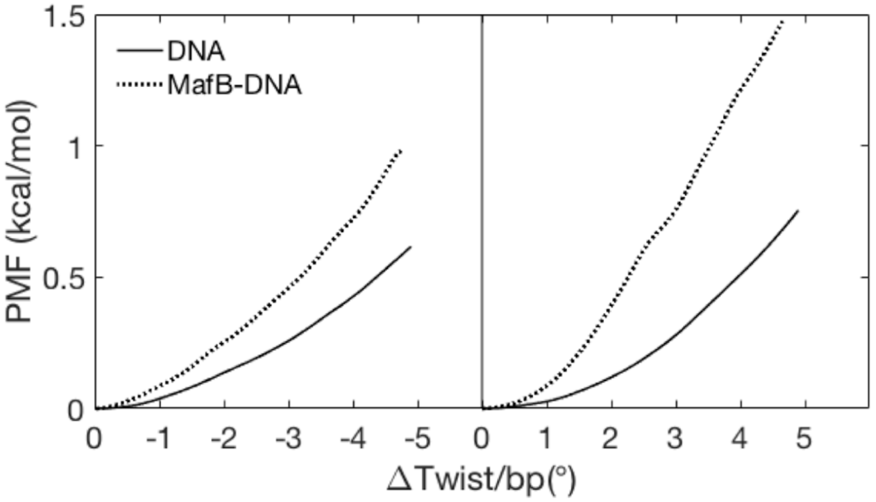


B

| | DNA | MafB-DNA |
|---|---|---|
| Relaxed tw (°) | 34.6 | 34.0 |
| K (kcal/mol*deg$^2$) | 0.057 | 0.11 |
| K+ | 0.069 | 0.11 |
| K- | 0.039 | 0.055 |
| C (pN nm$^2$) | 442 | 853 |
| P (nm) | 107 | 207 |

***Figure S3.*** **A).** PMF profiles for underwinding (**A**, left panel) and overwinding (**A**, right panel) of free DNA (black lines) and MafB-DNA (dotted lines) as a function of ΔTwist/b.p. **B)** Calculated average relaxed twists, torsional constants 'K' (overall), 'K-' (undertwisting regime), 'K+' (overtwisting regime), torsional moduli 'C', and torsional persistence lengths 'P' for MARE-DNA alone and in complex with MafB transcription factor. The force constants, obtained via quadratic regression, are used to calculate torsional moduli and the persistence lengths, as previously described[2].

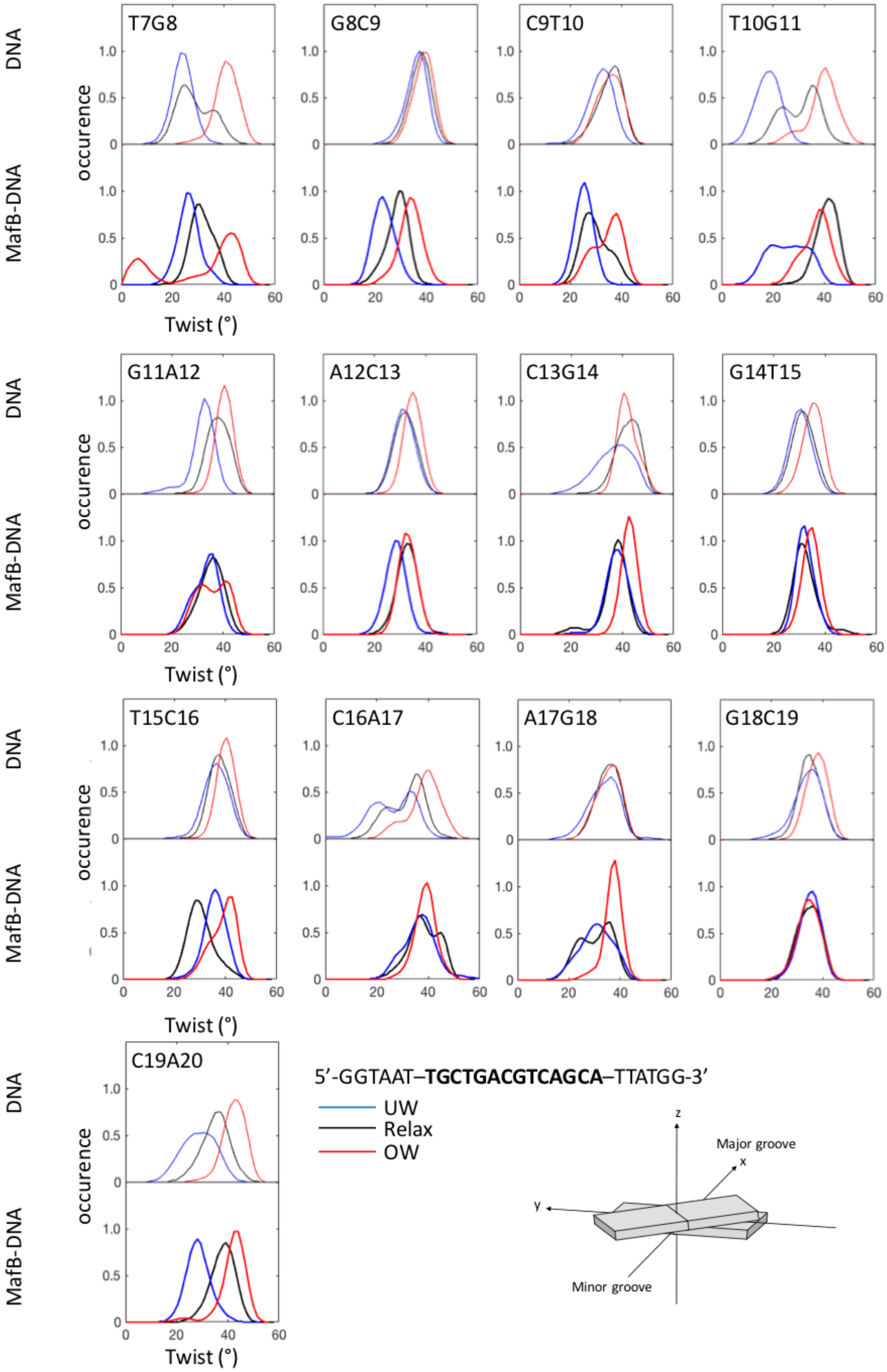


***Figure S4.*** Twist distributions of the restrained MARE region for underwound (-4.5), relaxed and overwound (+4.5) state. MafB-DNA is denoted with thicker lines.

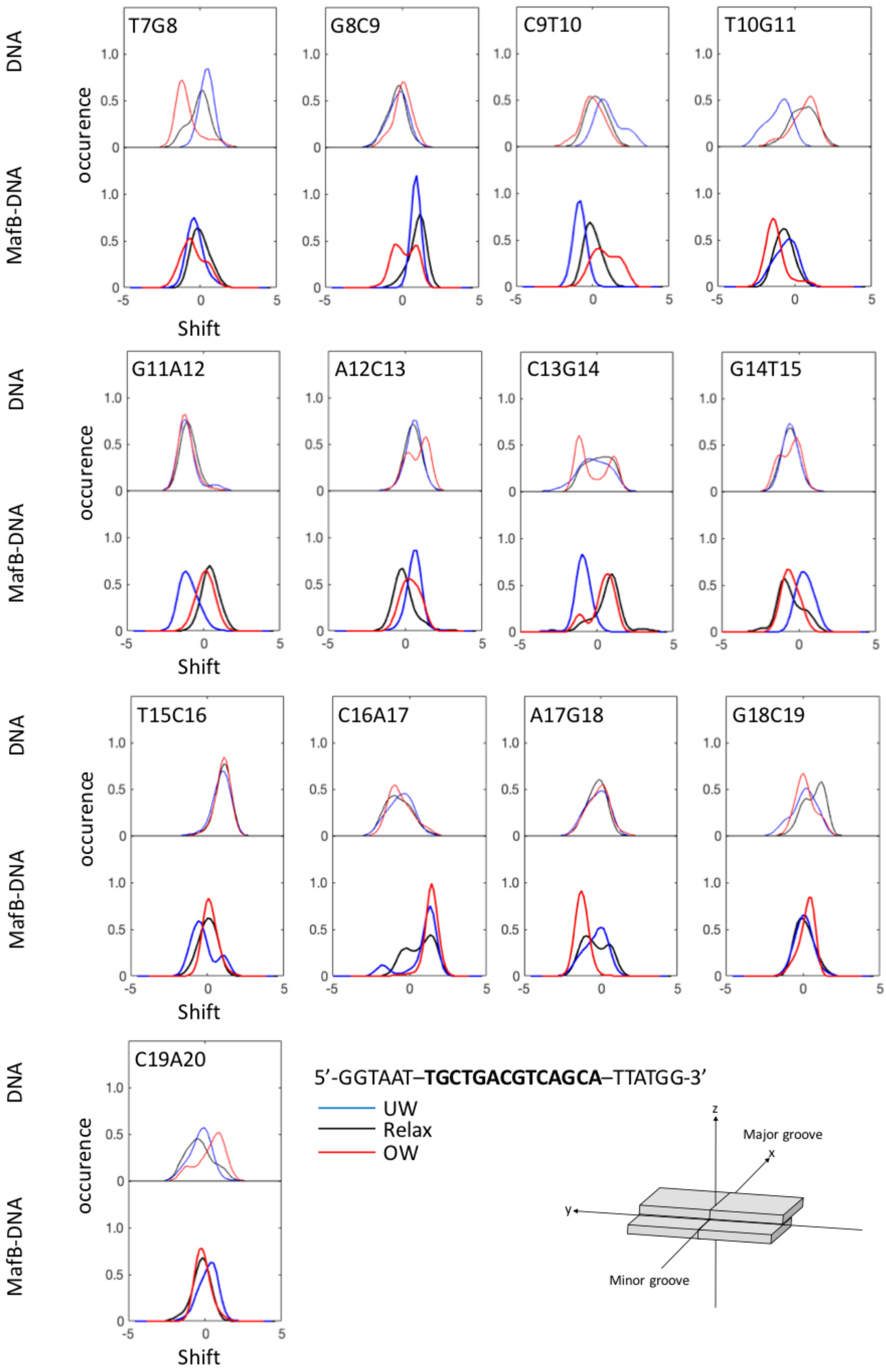


***Figure S5.*** Shift distributions of the restrained MARE region for underwound (-4.5), relaxed and overwound (+4.5) state. MafB-DNA is denoted with thicker lines.

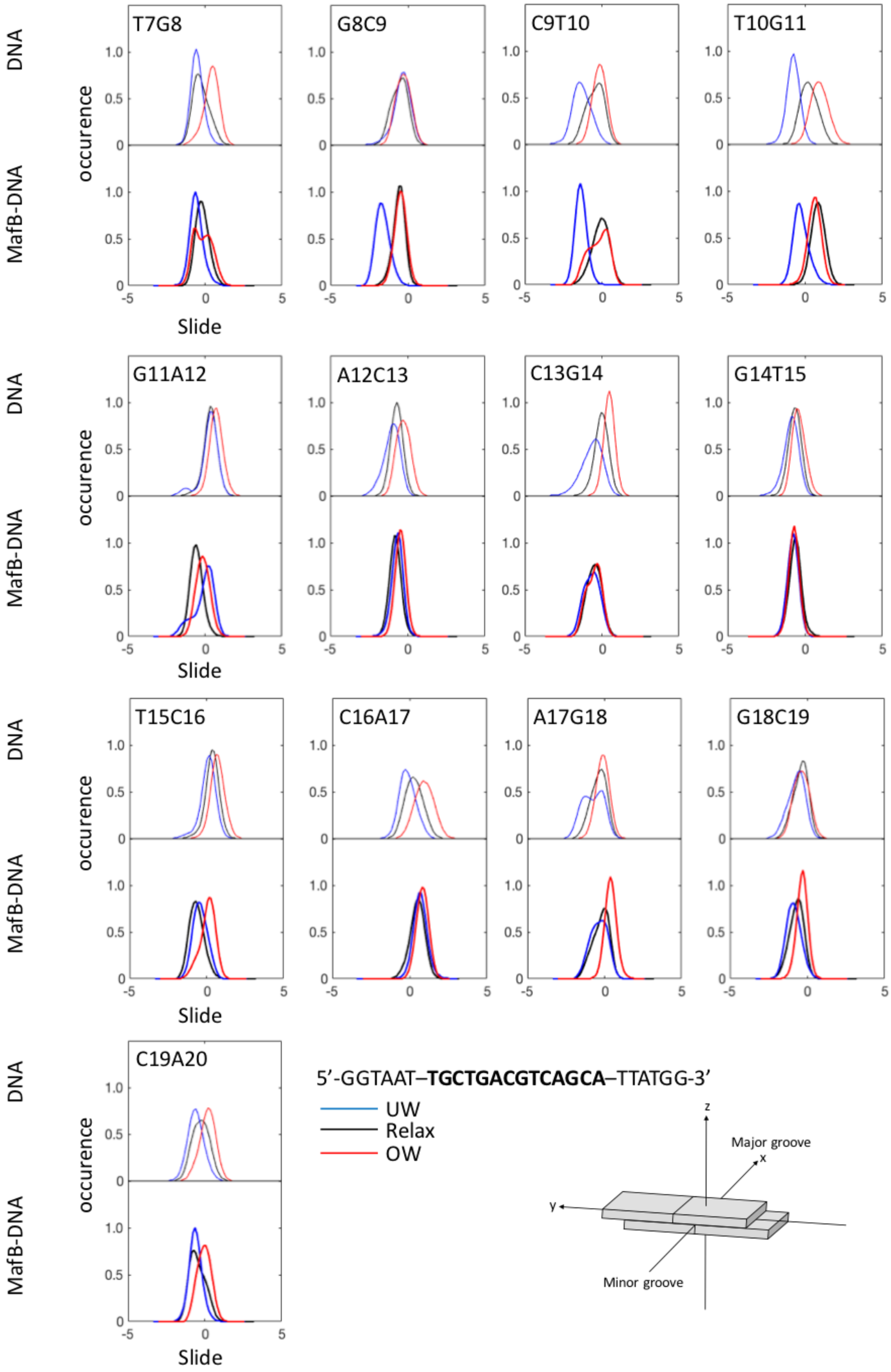


***Figure S6.*** Slide distributions of the restrained MARE region for underwound (-4.5), relaxed and overwound (+4.5) state. MafB-DNA is denoted with thicker lines.

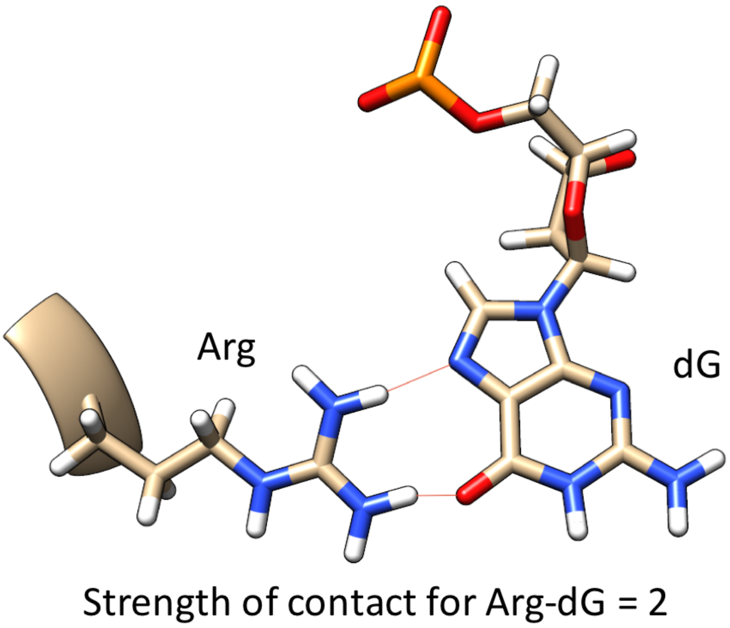


***Figure S7.*** The strength of MafB-DNA contacts is defined by summing all the interactions formed between the residues pairs. For example, an Arg-residue that forms two hydrogen bonds with a Gua-base has a contact strength of 2.

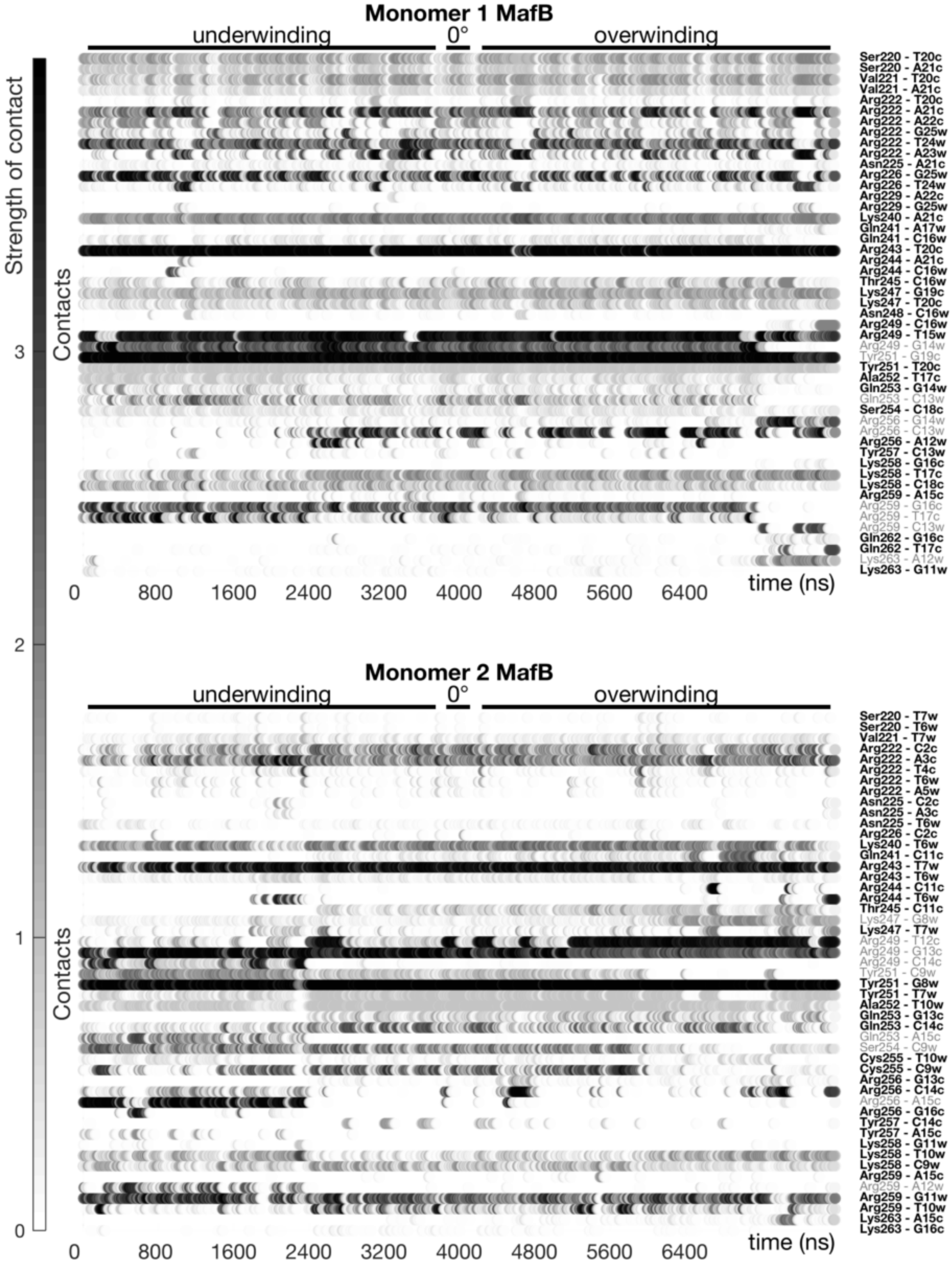


***Figure S8.*** Dynamic interactions map illustrating non-specific MafB-DNA contacts at different degrees of torsional stress. The interactions between pairs of the protein-DNA residues are characterized by a contact strength and occurrence. Torsional stress denoted as 'underwinding' represents changes from -5 degrees/b.p. to -0.5 degrees/b.p.; and 'overwinding' – from 0.5 degrees/b.p. to 5.0 degrees/b.p. Text in bold shows contacts that change insignificantly (change in contact strength < 1) with changing torsional stress. Indices "w" and "c" indicate to Watson- and Crick-DNA strands.

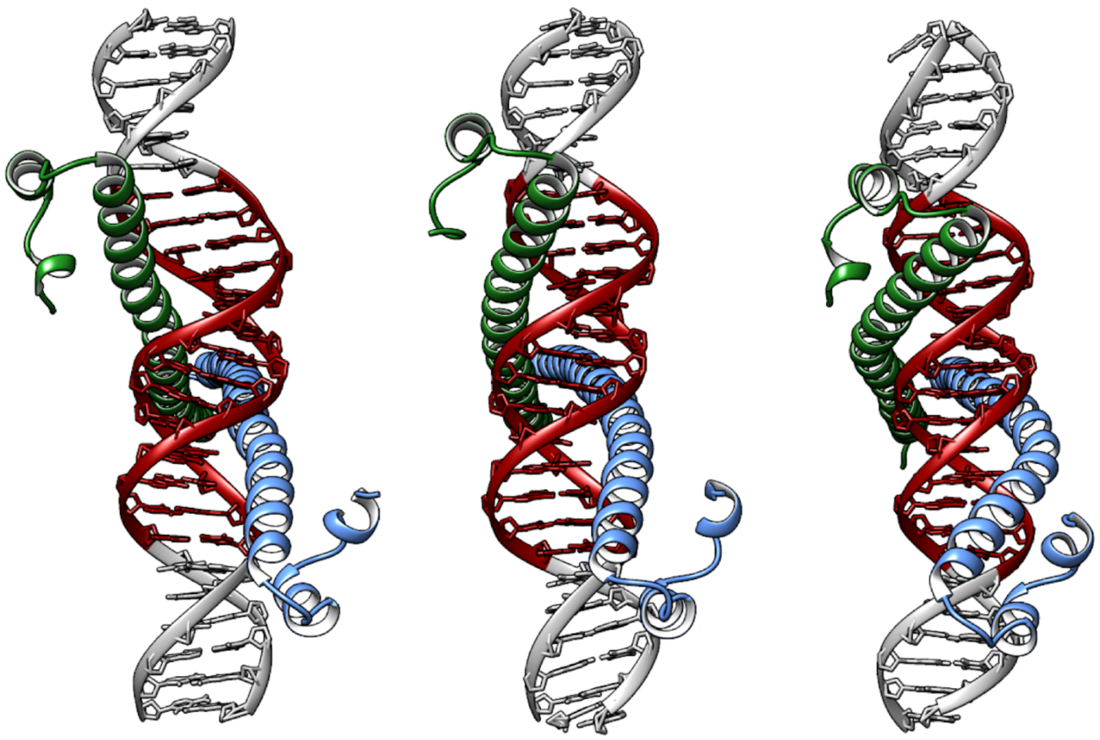


***Figure S9.*** Structural changes in the BZIP domain of the MafB-dimer at the underwound, relaxed and overwound states.
